# In vivo label-free observation of tumor-related blood vessels in small animals using a newly designed photoacoustic 3D imaging system

**DOI:** 10.1101/2021.11.09.467878

**Authors:** Yasufumi Asao, Kenichi Nagae, Koichi Miyasaka, Hiroyuki Sekiguchi, Sadakazu Aiso, Shigeaki Watanabe, Marika Sato, Shinae Kizaka-Kondoh, Yukari Nakajima, Kazuo Kishi, Takayuki Yagi

## Abstract

**Introduction:** Photoacoustic technology can be used for non-invasive imaging of blood vessels. In this paper, we report on our prototype photoacoustic imaging system with a newly designed ultrasound sensor and its visualization performance of microvascular in animal.

**Methods:** We fabricated an experimental system for animals using a high-frequency sensor. The system has two modes: still image mode by wide scanning and moving image mode by small rotation of sensor array. Optical test target, euthanized mice and rats, and live mice were used as objects.

**Results:** The results of optical test target showed that the spatial resolution was about 2 times higher than that of our conventional prototype. The image performance in vivo was evaluated in euthanized healthy mice and rats, allowing visualization of detailed blood vessels in the liver and kidneys. In tumor-bearing mice, different results of vascular induction were shown depending on the type of tumor and the method of transplantation. By utilizing the video imaging function, we were able to observe the movement of blood vessels around the tumor.

**Conclusion:** We have demonstrated the feasibility of the system as a less invasive animal experimental device, as it can acquire vascular images in animals in a non-contrast and non-invasive manner.

## INTRODUCTION

Animal imaging systems have become indispensable in basic research to elucidate biological activities and in the development of drugs. Conventional imaging devices for animal experiments include ultrasound, computed tomography (CT) and magnetic resonance imaging (MRI). To image blood vessels using these modalities, CT and MRI require a contrast agent. Ultrasound Doppler can image blood vessels without contrast, but the issue is that it depends on the skill of the surgeon, whether the object is an animal or a human.

On the other hand, photoacoustic technology has begun to be applied to devices for animal experiments. This technology is based on the imaging of photoacoustic waves generated by the light absorption of pigments, mainly hemoglobin, in living organisms, and it is possible to visualize vascular images non-invasively. The authors have developed a photoacoustic imaging system using a hemispherical detector array^1)^ (HDA) and have been promoting clinical research using this system in collaboration with several university hospitals and have demonstrated high-definition imaging of blood vessels^2)^. The feature of this system is that photoacoustic waves generated by light irradiation are received and reconstructed in almost all directions using an HDA. In other words, it is possible to construct a 3D volume image with a single pulse of light irradiation. In addition, by employing a scanning mechanism, it is possible to obtain a high-quality 3D image due to the averaging effect of scanning in wide area. This feature can also be used to generate 3D moving images with high reproducibility (so-called 4D imaging). In this report, we show the results of the evaluation of the newly designed HDA performance and also report the imaging performance of tumor-bearing mice to observe the changes in tumor-associated blood vessels induced by different cancer types.

## MATERIALS AND METHODS

### Device configuration

The photoacoustic imaging system (LUB-0) used in this report is a system developed by Luxonus Inc., that inherited the technology of the Japanese Cabinet Office program ImPACT^3)^. The LUB-0 system consists of a scanning stage with an HDA in the bed unit, a light source, optical fibers, a data acquisition system (DAS), and a personal computer (PC). In this paper, the horizontal plane is defined as the x-y plane, and the x-axis and y-axis are the short axis and long axis of the bed unit, respectively. The z-axis is the vertical axis (Supplementary Figure 1).

As in conventional devices such as the PAI-05, pulsed laser light is guided by an optical fiber and emitted upward along the z-axis from the bottom of the HDA, and a lens installed at the bottom of the HDA is used to spread the light in a conical shape to achieve a safe light irradiation density for the skin. The system can irradiate at two wavelengths, 797 nm and 835 nm, using either an alternating irradiation mode in which the two wavelengths are oscillated at 15 Hz and different wavelengths are alternately irradiated every 33 ms, or a single wavelength mode in which only one of the wavelengths is irradiated at 30 Hz. These can be selected on the user interface of operating monitor. In this experiment, all the images were taken using a single wavelength mode of 797 nm with 30 Hz laser irradiation.

As in the PAI-05, the objects were placed on a holding tray with a flat bottom on the bed unit for imaging. To maintain the flat bottom, a plastic mesh is used to support the object, and a sheet of polyethylene terephthalate (PET) film is used on the mesh to separate the acoustically matching water on the HDA from the water in the tray so that the object can be immersed in the water (Supplementary Video 1). In PAI-05, only the method of filling the tray with water was used for acoustic matching between the object on the tray and the HDA. On the other hand, in the LUB-0 system, ultrasound gel and other acoustic matching methods used in medical ultrasound systems can be employed.

For PA signal detection, a 512-channel receive-only transducer element was employed. The HDA for PA signal detection has a radius of 60 mm. The appearance of the sensor was almost the same as that of PAI-05. The differences in the specifications of the sensors used in LUB-0 and PAI-05 are shown in Table 1.

**Table 1.** Differences between the sensors used in this report (LUB-0) and that used in the previous report (PAI-05).

|  | Conventional (PAI-05) | New (LUB-0) |
| --- | --- | --- |
| Number of channels | 1024 | 512 |
| Radius of HDA | 55 mm | 60 mm |
| Center frequency | 3.34 MHz | 5.28 MHz |
| Bandwidth | 85 % | 81 % |

The same receiving frequency of the sensor as that of PAI-05 was prototyped as a comparative example, and a new sensor was designed and prototyped for the animals reported here, with the center frequency of the sensor shifted to the high-frequency side to enable reception of higher-frequency signals. Along with this change in the sensor, the electrical circuit of the receiving signal processing system was also improved. Although the number of channels has been halved compared to PAI-05, the effect of the improved averaging frequency due to the 1.5 times faster laser repetition rate confirms that the design image quality is equivalent for the same received frequency band.

### Data acquisition

The PA signals generated by pulsed laser light irradiation from the laser unit were received by the HDA of 512 channels and the signal data were transferred to the DAS, which sampled PA signals at 60 MHz. The sampled digital data was sequentially transferred to the PC for image reconstruction.

Image reconstruction was performed in real time during scanning, and the reconstructed PA images were displayed on the operation PC.

Video images from the digital camera installed at the bottom of the HDA were displayed on the screen during scanning simultaneously with the transfer of PA signals. The acquired camera images were combined during scanning to produce a photographic image of the entire scanning area.

As with the PAI-05 system, two types of PA images are available: the “STILL mode” in which the HDA scans and captures a wide area, and the “MOVIE mode” in which the sensor is fixed or moved slightly to a specific location and the averaged PA image of that location is updated in real time.

In the still image mode, the scanning area of the HDA was expanded by drawing a rectangular orbit and gradually increasing the length of each side from the center. In addition to this, a raster scan method with a slower scanning speed is adopted, and a method to acquire images with higher SN by increasing the number of overlaps is adopted. These two scan sequences can be selected by the user. In LUB-0, any rectangular size can be selected within the maximum measurement area of 290mm × 180mm or less.

The movie mode allows two types of HDA scan operations: one is called simple movie (SM) mode, in which the HDA remains stationary on a spot and repeatedly irradiates a laser beam on one spot to update the PA image. The other is called the fluctuation motion (FM) mode, in which the image is captured in a continuous micro rotational. In both movie modes, temporal images for reconstructed images can be accumulated and averaged according to the number of times the laser beam is irradiated. In the FM mode, the scanning motion was performed while the HDA was rotated so that its center followed the path of a circle with a diameter of 3 mm. Furthermore, in movie mode, the speed of sound, which is a parameter for image reconstruction, can be changed while the image is being captured. This allows the user to set the optimal sound speed value by adjusting the focus while viewing the reconstructed image.

### Imaging method

Universal Back Projection^4)^ (UBP) was used to reconstruct the PA images. The default value of the reconstructed voxel size was 0.1 mm in each xyz direction, and the voxel size could be changed according to the size of the object and the purpose of analysis. In addition, the sound speed for reconstruction can be changed to allow the user to adjust to the optimum sound speed or to obtain the more focused image. A newly developed program (PAT viewer) was used for image display.

### Animal experiments

All animal experiments were performed with the approval of the respective animal experiment ethics committees of Tokyo Institute of Technology and Keio University. In addition, human subjects were scanned in accordance with the internal ethics regulations of Luxonus Inc. and managed to prevent the identification of the individual. Male BALB/c nude mice were purchased from Oriental Yeast Co., Ltd. (Tokyo, Japan). Male HWY/Slc hairless rats were purchased from Sankyo Labo Service Corporation, Inc. Male Jcl:SD rats were purchased from CLEA Japan, Inc. All mice and rats used were 5–9 weeks of age.

### Imaging subjects

The phantom used in this study was the USAF1951 chart^5)^, which is used for performance evaluation in the field of optics. The chart is patterned by chromium metal on a glass substrate, with a pattern of several groups of six elements of different sizes, where each element consists of three lines and spaces. The correspondence table of each element and line and space is shown in Supplementary Table-1.

Euthanized nude mice, hairless rats, and SD rats were used as the phantoms closest to the living bodies. As a method of euthanasia, a CO2 euthanasia method was adopted instead of cervical dissection with trauma to prevent damage to blood vessels. The image of a healthy male subject was used as the subject for the image evaluation test using a living body.

Images of tumor-bearing mice were taken using several tumor models. The tumor cells used were mouse breast cancer cell line 4T1 and mouse osteosarcoma cell line LM8. For 4T1, in addition to orthotopic transplantation into the right fourth mammary gland, we used a subcutaneous model in which small pieces of tumor block from another mouse used for imaging were transplanted to the back. For LM8, we used a subcutaneous transplant model on the thigh.

## RESULTS

### Comparison with conventional sensors - 1 - Phantom experiment

The performance of the system was evaluated using the USAF1951 chart phantom. The results are shown in Fig. 1(a)-(d). Figure 1(a) shows an image taken by an HDA with conventional performance, which is a comparative example. Figure 1(b) is the image obtained by the newly designed HDA, and Fig. 1(c) is a magnified view of the dashed square in (b). With the conventional HDA, the element indicated by the blue triangle in Fig. 1(a) was visually separated. This means that a 0.177 mm wide line and space can be resolved. In the new HDA, the element shown by the yellow triangle in Fig. 1(c) was visually separated. Figure 1(d) shows a graph of the luminance profile on the lines indicated by A and B in Fig. 1(c). This means that a 0.099 mm wide line and space can be resolved. This demonstrates that the device using the new HDA has a spatial resolution of less than 0.1 mm.

**Figure 1.**
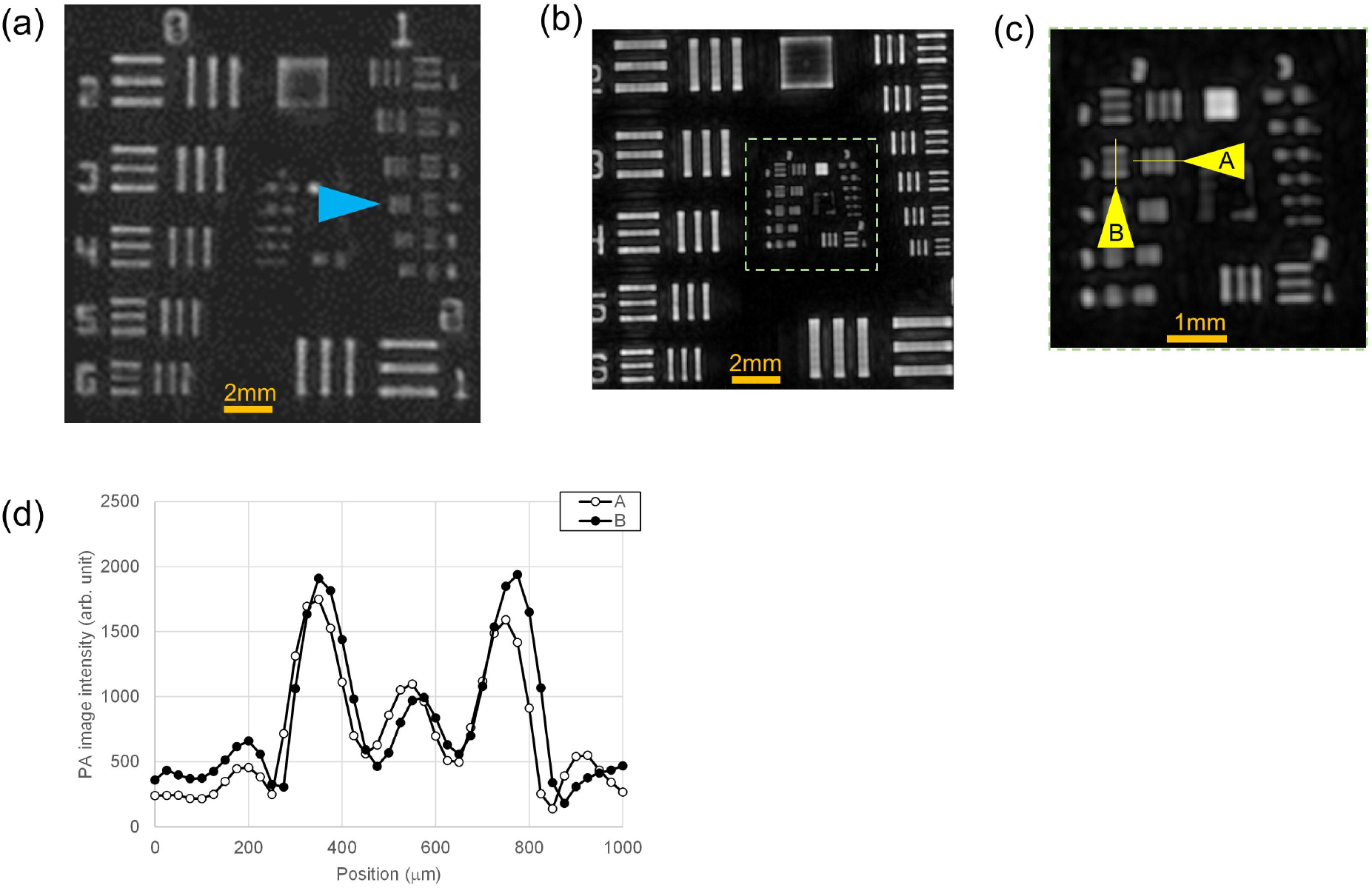
The performance of LUB-0 system evaluated using the USAF1951 chart phantom. (a) PA image taken by an HDA with conventional performance, which is a comparative example. (b) PA image obtained by the newly designed HDA. (c) The magnified view of the dashed square in (b).

### Comparison with conventional sensor-2 - Human body

Next, to compare the performance of the new HDA and the conventional one, we acquired images of the ring finger of the same person using different devices. The images were taken on different days. The results are shown in Fig. 2(a)-(b). Figure 2(a) shows the image taken by the HDA with conventional performance, which is a comparative example. Figure 2(b) is the image obtained by the newly designed HDA. This comparison shows that the new HDA visualizes more blood vessels and even peripheral branches are imaged.

**Figure 2.**
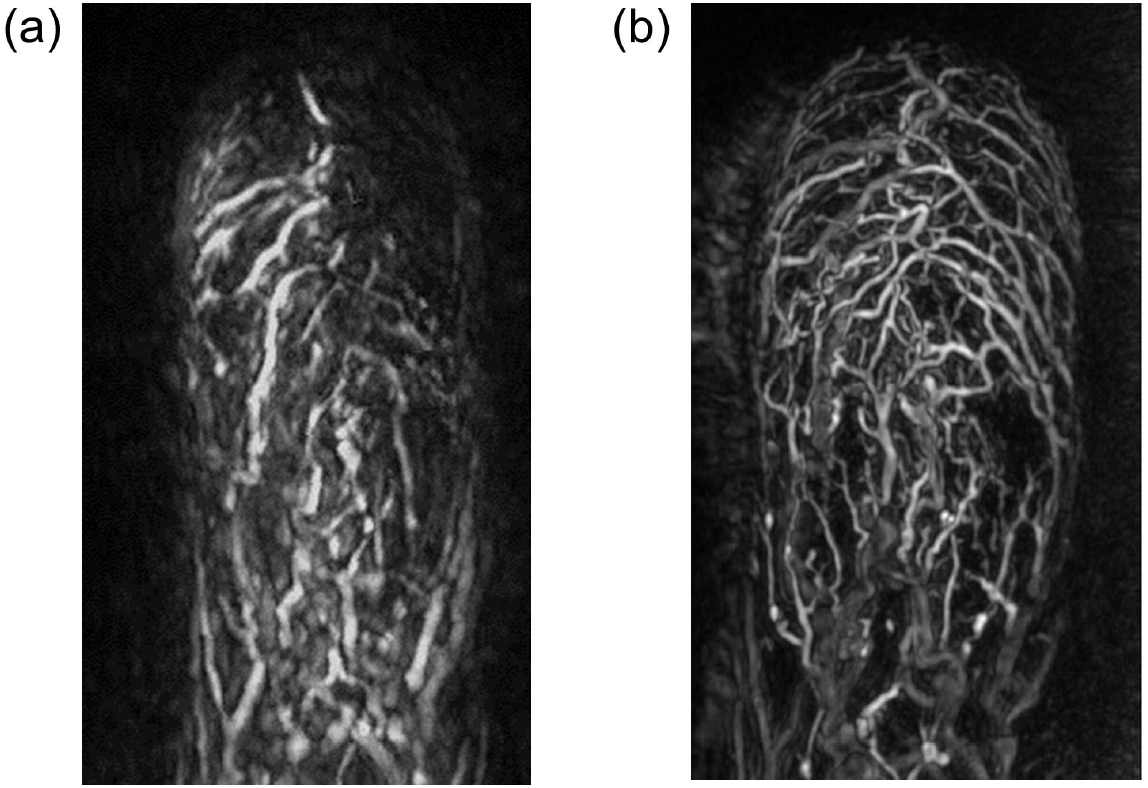
PA images of the ring finger of the same person using different devices. (a) PA image taken by the HDA with conventional performance, which is a comparative example. (b) PA image obtained by the newly designed HDA.

### Imaging performance of small animals

Images were acquired after euthanasia of small animals that had not been subjected to prior intervention by other experiments. The results for nude mice, hairless rats, and SD rats are shown in Figures 3 through 5.

**Figure 3.**
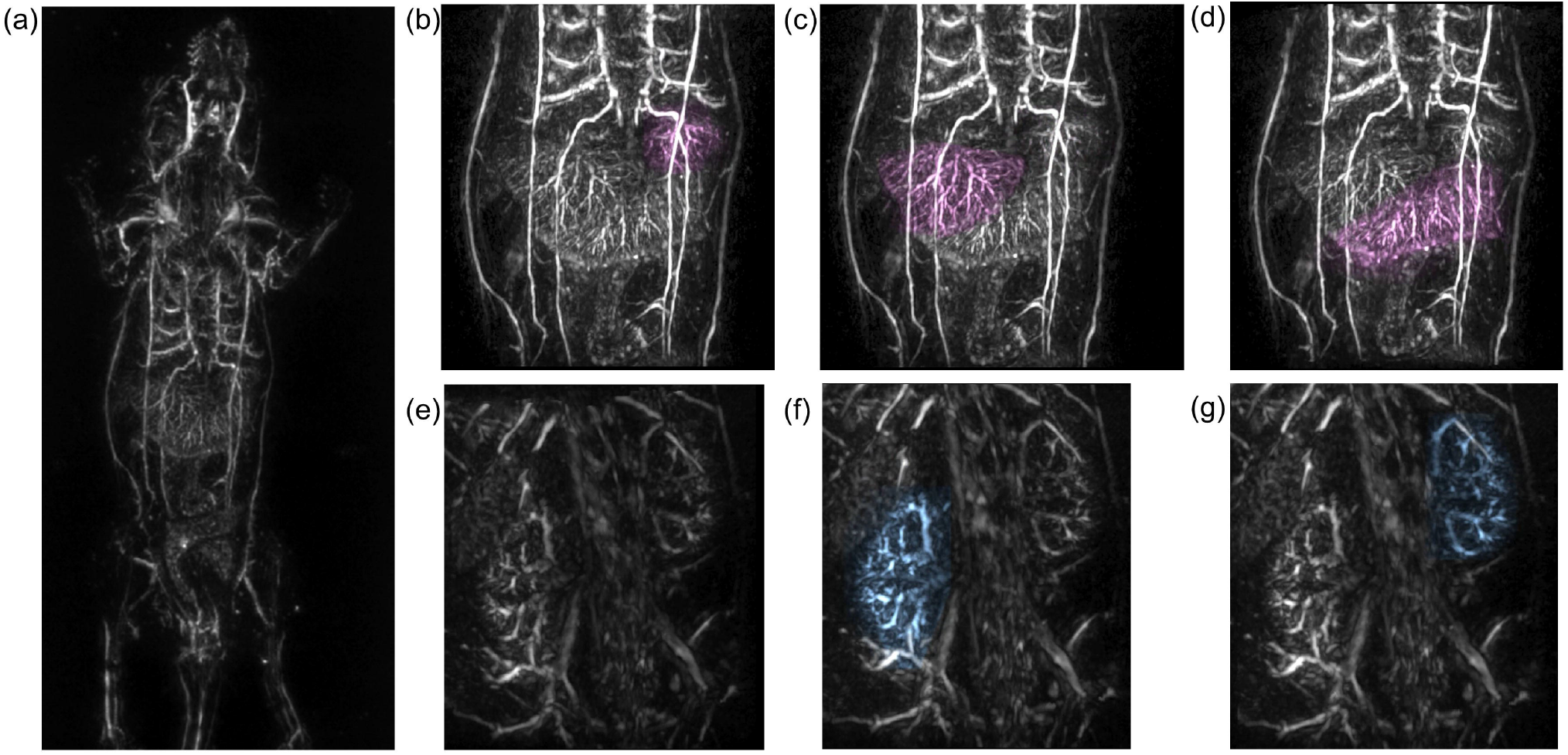
PA image of the whole body of a nude mouse. (a) The vascular image of the whole body taken in the prone position. (b)-(d) Magnified PA images of the liver in the image in (a), with each lobular structure manually highlighted using the viewer’s functions. (e) The magnified image of the back of a nude mouse in the supine position. (f)-(g) PA images of right and left kidneys in (e) manually highlighted using the viewer’s functions.

Figure 3(a) shows a photoacoustic image of the whole body of a nude mouse taken in the prone position. The vascular image of the whole body is depicted. Especially in the liver, we can observe the vascular branching based on multiple lobular structures. Figure 3(b)-(d) shows the results of manual coloring of each of these lobular structures using the newly developed viewer functions (Supplementary Video 2). Figure 3(e) shows a magnified image of the back of a nude mouse in the supine position. The results of manually coloring the left and right kidney regions are shown in Figure 3(f)-(g). In this way, the aorta, renal vessel, arch vessels in the kidney, and the branching vessels toward the nephron area were visualized (Supplementary Video 3). Figure 4 (a) shows an image of a hairless rat in the prone position and (b) shows an image of a hairless rat in the supine position. Similar to nude mice, images of the liver and kidneys and other organs are clearly visualized. From Fig. 4(b), it was possible to visualize the vascular structure of the brain through the skull (Supplementary Video 4).

**Figure 4.**
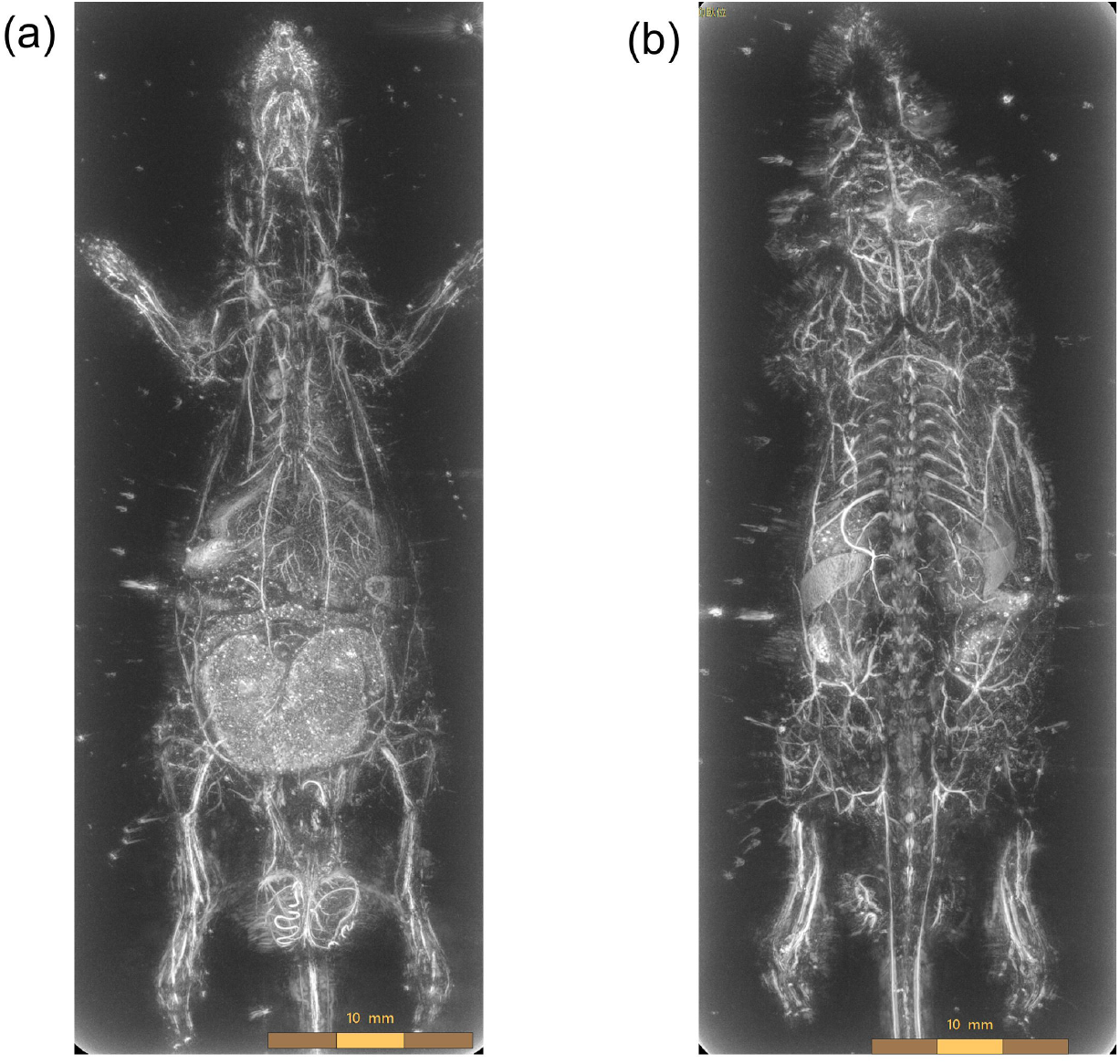
PA images of a hairless rat. (a) Prone position (b) Supine position.

Scans were also performed on SD rats, which are larger in size than hairless rats. These results, together with images of human palms and legs, are shown in Fig. 5, which shows an example of objects of various sizes.

**Figure 5.**
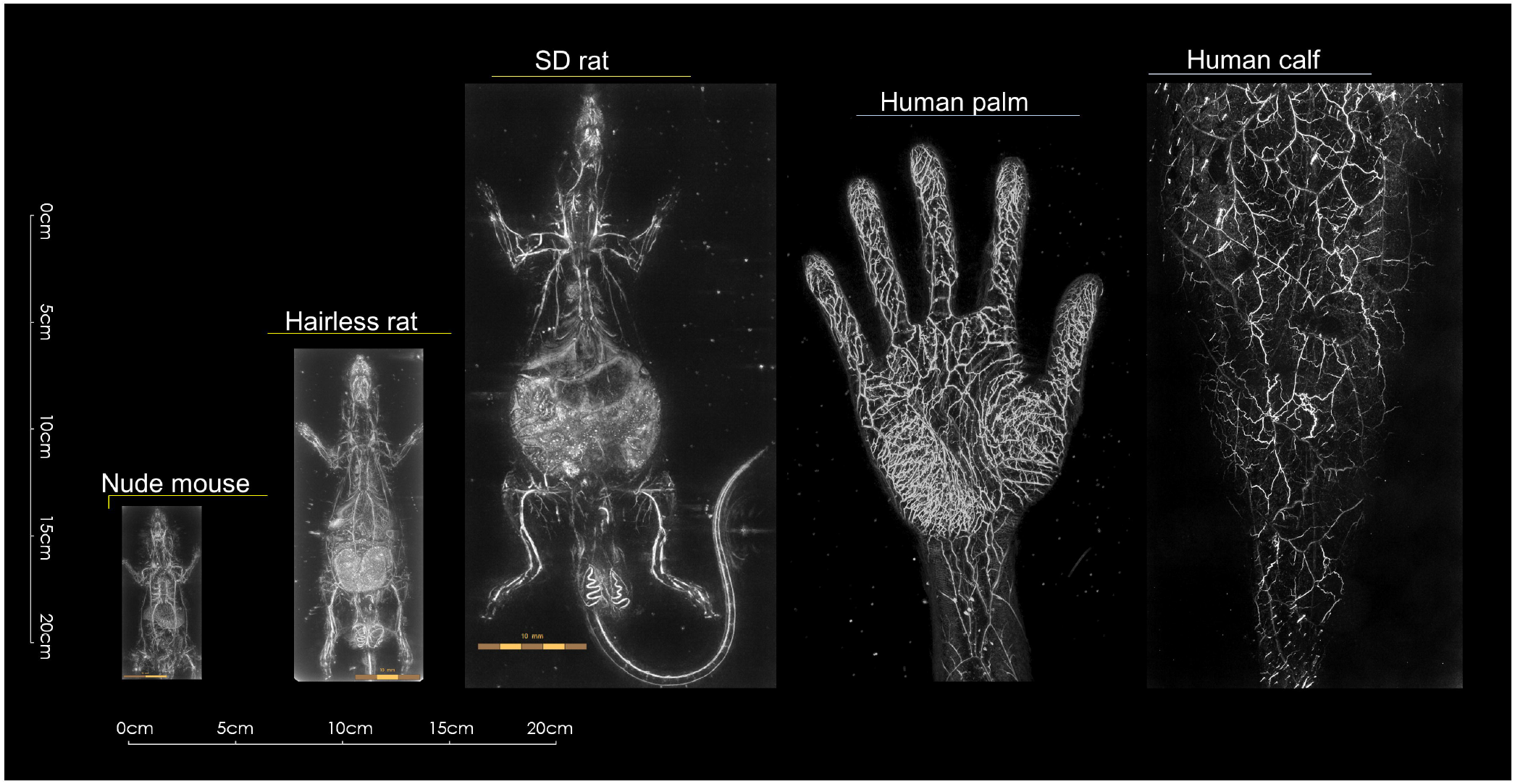
An examples of PA images taken with objects of various sizes, including a nude mouse, a hairless rat, an SD rat, a human palm, and a human calf.

Next, images of tumor-bearing mice with two types of tumors, breast tumor and osteosarcoma, were taken: euthanized mice and mice that were scanned under anesthesia while still alive. Figure 6(a)-(c) show the images of mice with LM8 implanted on the thigh and allowed to grow in a supine position after euthanasia. (a) is a whole image, (b) is a magnified image of the tumor area, and (c) is a partially cropped image to make it easier to see the inside of the tumor. The signal from the edge of the tumor is very strong, suggesting that the tumor is rich in blood flow, but the partially cropped image suggests that there is little signal from inside the tumor.

**Figure 6.**
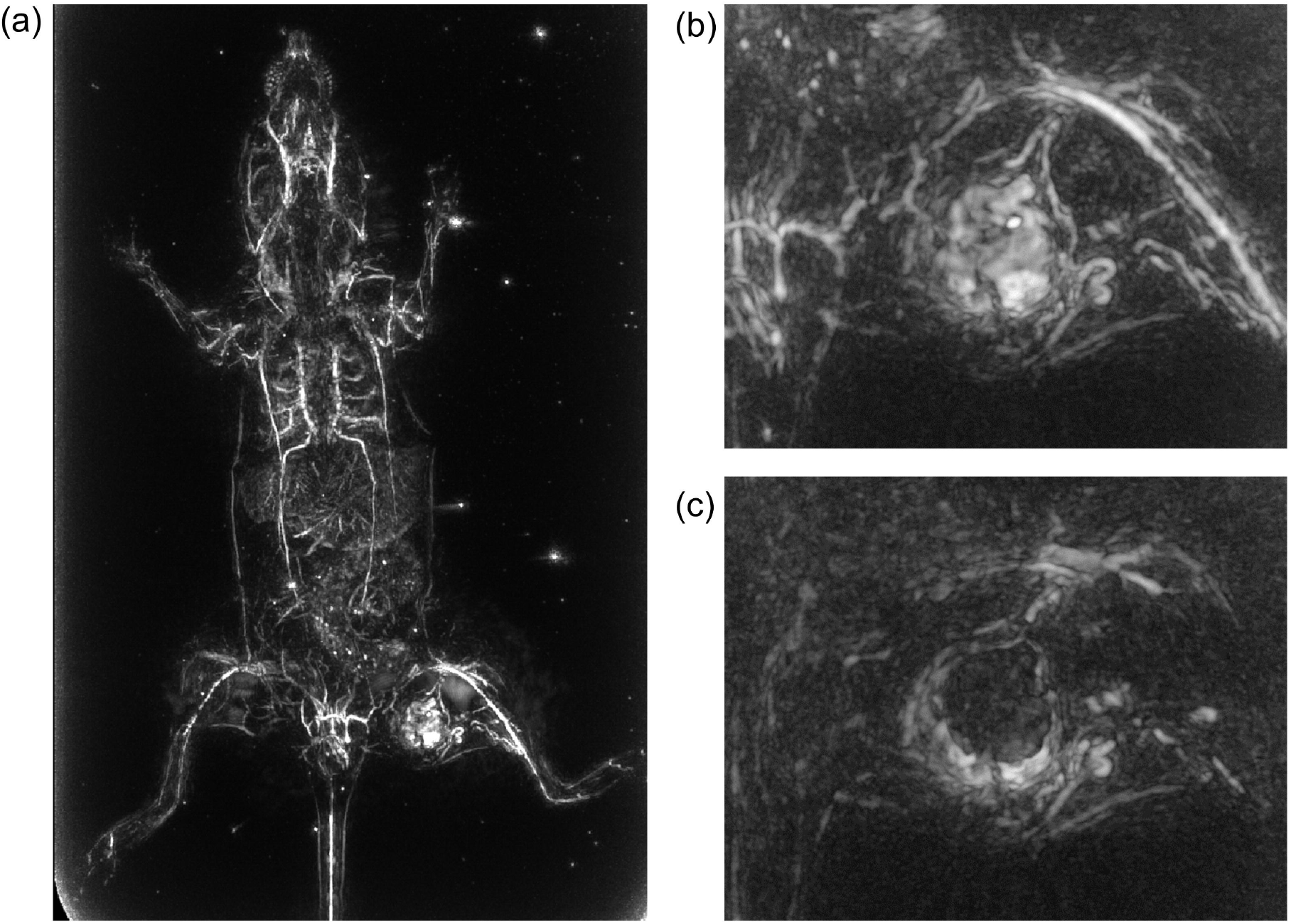
PA images of a mouse with sarcoma-derived tumor cells LM8 implanted in the thigh and allowed to grow in a supine position after euthanasia. (a) A whole body image. (b) A magnified image of the tumor area. (c) A partially cropped image to make it easier to see the inside of the tumor.

Figure 7 (a)-(b) show images of orthotopic 4T1 tumor-bearing mice in the prone position after euthanasia. Figure 7(a) is the image below the chest and Fig. 7(b) is a magnified image of the tumor area. The signal from the tumor edge was absent, suggesting the induction of blood vessels toward the tumor. Figure 7(b) is a brightened image by adjusting the image window level. The signal from the inside of the tumor was relatively diffuse, suggesting that blood flow also existed inside the tumor. Figure 8 (a)-(e) show images taken in the supine position of the mouse with a subcutaneous 4T1 tumor, which was formed after small pieces of tumors block of 4 mm in size were transplanted to the back from an orthotopic tumor about 15 mm in diameter in another mouse. The mouse was alive and under inhalation anesthesia. Figure 8(a) is the whole image. When observed together with the composite image from the digital camera embedded in the HDA, it is possible to see how many blood vessels are guided toward the location of the tumor mass (Supplementary Video 5). Fig. 8(b) is an enlarged image of the tumor area, and Fig. 8 (c) is a partially cropped image. The results suggest that blood vessels to the tumor are heavily induced, and blood vessels surrounding the tumor area are clear, but there is little signal from inside the tumor in the cross-sectional image. Video imaging was performed on these tumor images. We used the FM mode and averaged the images 9 times. As a result, the movement of blood vessels in the depths was observed. The yellow arrowheads in Figure 8(d)-(e) indicate that the blood vessels are moving. This was observed to fluctuate with a period of about 2.5 seconds (Supplementary Video 6). On the other hand, the signal from the inside of the tumor showed many fine blood vessels compared to the static cross-sectional image.

**Figure 7.**
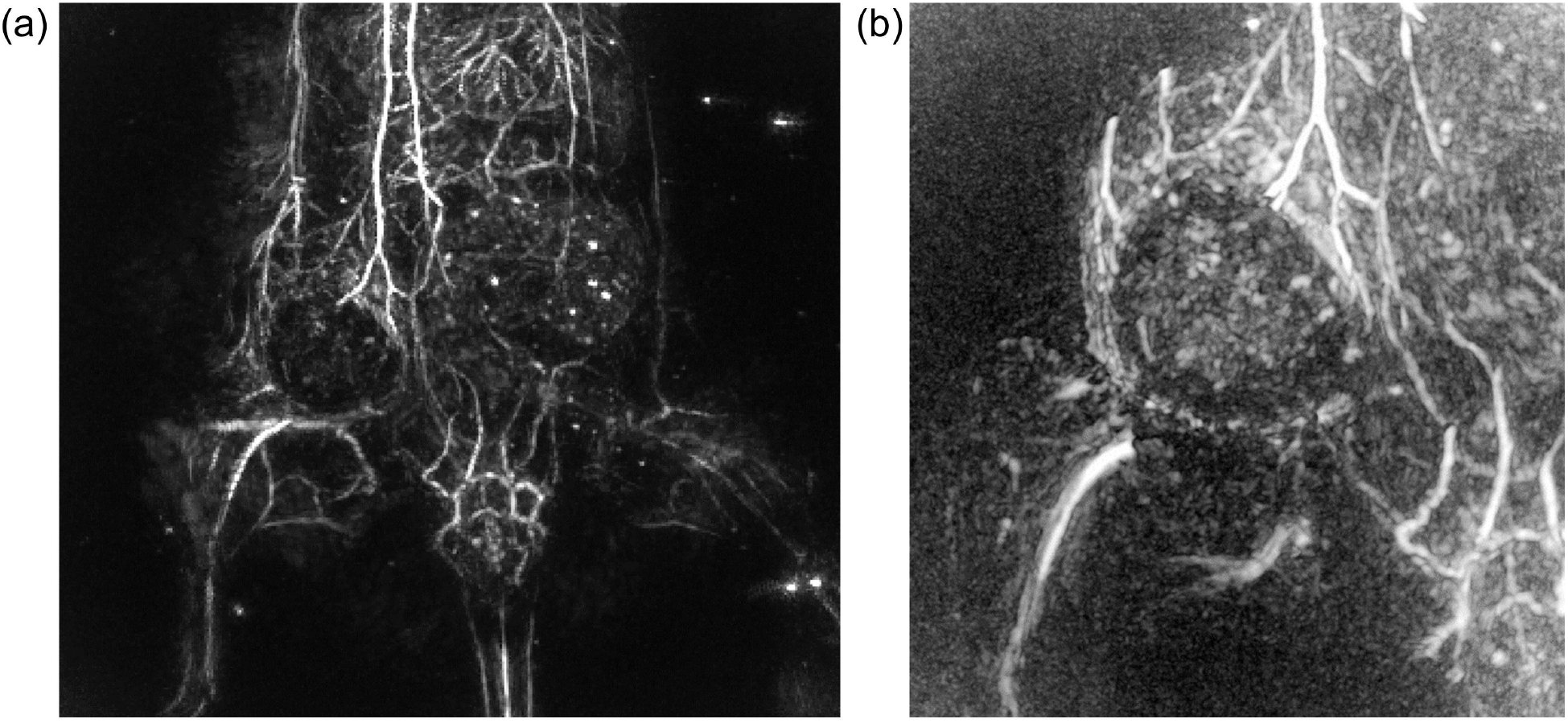
PA images of a mouse in the prone position after euthanasia, in which 4T1 tumor cells derived from breast cancer were implanted into the right fourth mammary gland and allowed to grow. (a) PA image below the chest. (b) A magnified image of the tumor area.

**Figure 8.**
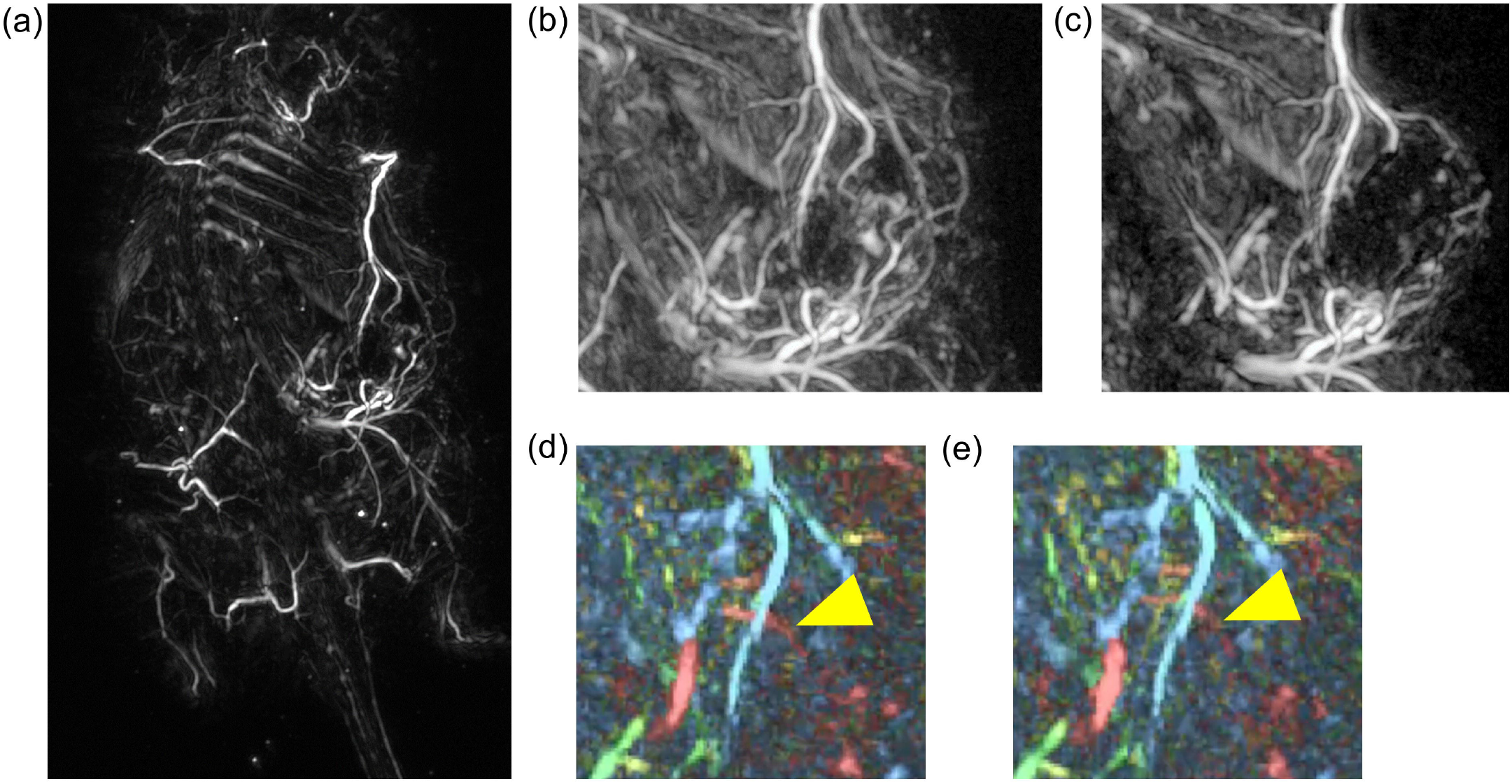
PA images taken in the supine position of a mouse in which 4T1 tumor cells derived from breast cancer were implanted in the right fourth mammary gland of another mouse, the tumor was grown to a diameter of about 15 mm, removed and divided into blocks of 4 mm in size, and the blocks were implanted in the back to grow the tumor. The mouse was alive and under inhalation anesthesia. (a) The whole-body image, (b) An enlarged image of the tumor area. (c) A partially cropped image. (d)-(e) PA images indicating that the blood vessels are moving.

The results obtained in this study are shown in Table 2.

**Table 2.** Summary of the results of induced tumor vascularity in tumor-bearing mice obtained in this report.

|  | Sarcoma | Breast cancer-1 | Breast cancer-2 |
| --- | --- | --- | --- |
| Cell line | LM8 | 4T1 | 4T1 |
| State during scanning | post-euthanasia | post-euthanasia | Alive, under anesthesia |
| Transplant Materials | Cell | Cell | Tumor block |
| Vascular induction | ✓ ✓ ✓ | ✓ ✓ | ✓ ✓ ✓ |
| Tumor edge signal intensity | ✓ ✓ ✓ | ✓ | ✓ |
| Vessel-like signals inside the tumor | ✓ | ✓ ✓ | Still ✓<br>Movie ✓ ✓ |

## DISCUSSIONS

By using an optical test target, we demonstrated that a new type of sensor capable of receiving high frequency signals can achieve a spatial resolution about twice that of the conventional prototype. At the same time, we demonstrated with an image of a human fingertip that it is possible to visualize minute vascular branching structures that could not be resolved in the past. In general, there is a trade-off between resolution and depth of field in diagnostic devices that use the ultrasound principle. We believe that this new sensor-based system is no exception to this, but we did not evaluate the depth performance in this experiment, which is an issue for the future. Nevertheless, in this experiment, organs relatively close to the subcutaneous tissue such as the liver and kidneys were clearly visualized in small animals such as mice and rats. Other organs such as the spleen and digestive tract were also visualized. Therefore, it can be said that the depth performance is generally sufficient for basic research in animal experiments.

In experiments with tumor-bearing mice, different morphological vascular structures were induced depending on the tumor type and implantation site. It is already known that the induction of vascular structure differs depending on the type of tumor and the microenvironment, but our devise clearly visualized and confirmed the difference. In addition, it seems that not enough research has been done on the differences in the induction of blood vessels depending on the transplantation method. We look forward to further research to elucidate the mechanism of tumor neovascularization. Furthermore, we observed that the blood vessels in this tumor vascular structure move with the movement of the body. The video shows that the oscillation period is about 24 times per minute, which cannot be explained by the average heartbeat of mice (500 beats per minute) or the average respiratory rate (160 beats per minute), so it may be due to a different mechanism. Since the tumor on the back was scanned from the supine position, it may be caused by peristalsis of the gastrointestinal tract, but a detailed analysis of the mechanism is a future issue.

Although the verification of the usefulness of such 3D moving images, so-called 4D images, is expected in future research, it is considered to be difficult to realize with the other photoacoustic imaging systems that have been reported so far. In the acoustic resolution PA microscope^6)^ (AR-PAM), the sensor itself is moved to scan the acoustic acquisition position point by point using a focus transducer. Since the sensor itself has to be moved, it takes time to acquire one image, making video imaging difficult.

Optical resolution PA microscopy^6)^ (OR-PAM) can acquire images at video rates because light scanning can be accelerated by using mirror optics, but imaging of the entire tumor cannot be achieved because the imaging range is limited to the receivable range of a single ultrasonic sensor and imaging of deep areas is not possible.

The method of rotating the arc-shaped sensor around the object^7)^ allows 3D imaging over a wide range, but it takes time to rotate the sensor around the object, so video imaging is not possible. Similarly, the method of mechanically scanning the linear sensor in the direction of sensor elevation^8)^ is also unsuitable for 3D moving images. This is a strong point of the hemispherical sensor, which enables three-dimensional imaging with a single shot of laser irradiation as in this system. In addition, the scanning mechanism functions effectively to obtain a high-quality image.

As described above, the use of high-quality 3D still images and 3D moving images in the research field is expected to replace some of the animal experiments that have been conducted using conventional methods. In other words, for example, in tests to evaluate the effects of drugs, the evaluation of the effects on the organs of mice has conventionally had to rely on autopsy evaluation. It is also desirable from the viewpoint of animal welfare. In addition, this system can be used to measure both animals and humans using the same principle. This can be expected to contribute to seamless medical research.

In addition, it has already been demonstrated through clinical studies that the use of dyes such as indocyanine green (ICG) can achieve higher resolution imaging of lymphatic vessels than the fluorescence method^9)^. In the future, it is expected that basic research on molecular imaging using antigen-antibody reactions, as well as the use of fluorescent agents, can be realized based on higher resolution images by using this system.

## SUMMARY

As described above, we have completed a high-resolution PA imaging system that is approximately twice as large as conventional systems and has been able to produce images of various sizes of objects. The ability to image large animals means that the system can be used in a wider range of applications than the mouse dedicated CT or MRI. In addition, peripheral microvessels was depicted more clearly than before. By being able to observe what was previously invisible, it is expected to open up new research and clinical applications. It is also expected to be applied to the field of molecular imaging, which was not examined in this study. As described above, we have demonstrated the feasibility of this system for use in animal experiments through the introduction of various images, and we strongly hope that it will contribute to the development of medical technology by spreading the use of this system and promoting more detailed research in various fields.

## ACKNOWLEDGEMENTS

This development is supported by the Japan Agency for Medical Research and Development (AMED) and the New Energy and Industrial Technology Development Organization (NEDO).

## FIGURE LEGENDS

Supplementary Figure 1

The photoacoustic imaging system (LUB-0) used in this report. The horizontal plane is defined as the x-y plane, and the x-axis and y-axis are the short axis and long axis of the bed unit, respectively. The z-axis is the vertical axis.

Supplementary Table 1

Correspondence table between each element and line and space of the USAF 1951 chart used as a phantom in this report.

Supplementary Video 1

Video showing a small animal being placed on a holding tray for scanning by LUB-0 system.

Supplementary Video 2

PA image video of a nude mouse taken in the prone position.

Supplementary Video 3

PA image video of a nude mouse taken in the supine position.

Supplementary Video 4

PA image video of the brain blood vessel through the skull of a hairless rat.

Supplementary Video 5

Video with digital camera image and PA image superimposed.

Supplementary Video 6

Video showing vascular oscillations deep within the tumor in tumor-associated blood vessels of a tumor-bearing mouse.

