## Supplementary figures and images for "In vivo label-free observation of tumor-related blood vessels in small animals using a newly designed photoacoustic 3D imaging system"

### Supp. Figures

#### Slide 1
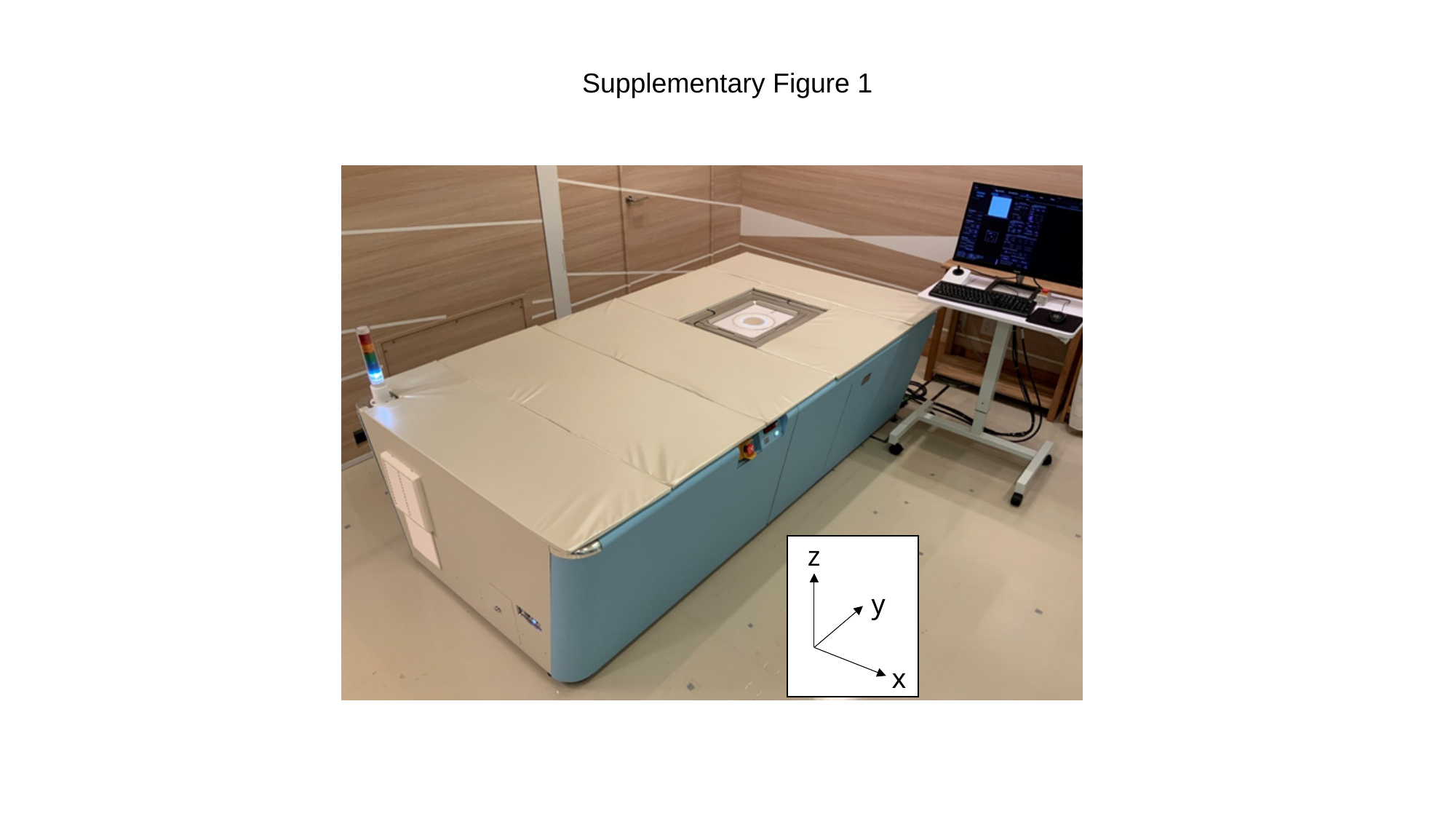

Supplementary Figure 1
z
y
x

#### Slide 2
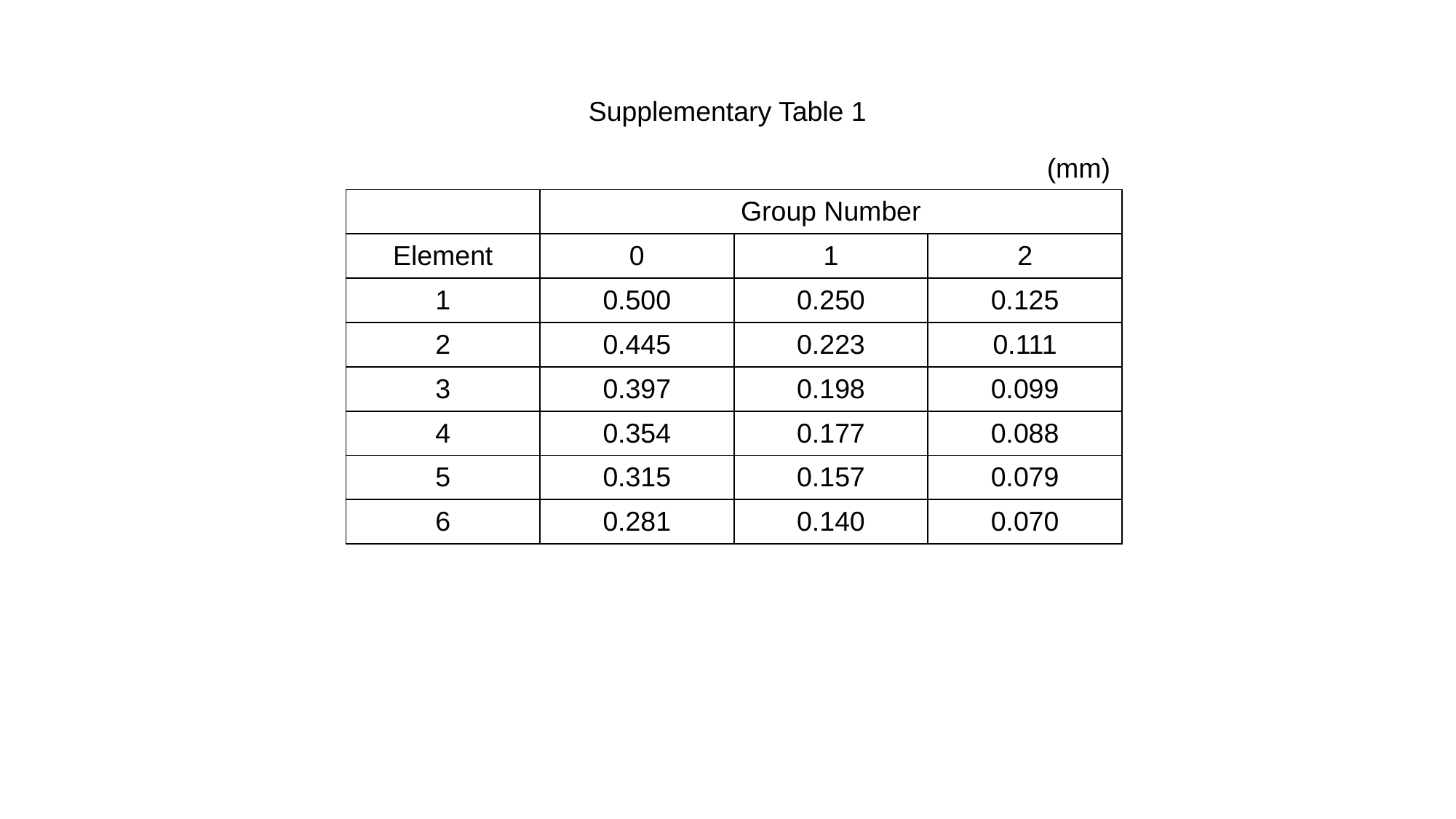

Supplementary Table 1
(mm)
| | Group Number | | |
| --- | --- | --- | --- |
| Element | 0 | 1 | 2 |
| 1 | 0.500 | 0.250 | 0.125 |
| 2 | 0.445 | 0.223 | 0.111 |
| 3 | 0.397 | 0.198 | 0.099 |
| 4 | 0.354 | 0.177 | 0.088 |
| 5 | 0.315 | 0.157 | 0.079 |
| 6 | 0.281 | 0.140 | 0.070 |
